# Mapping native R-loops genome-wide using a targeted nuclease approach

**DOI:** 10.1101/457226

**Authors:** Qingqing Yan, Emily J. Shields, Roberto Bonasio, Kavitha Sarma

## Abstract

R-loops are three-stranded DNA:RNA hybrids that are pervasive in the eukaryotic and prokaryotic genomes and have been implicated in a variety of nuclear processes, including transcription, replication, DNA repair, and chromosome segregation. While R-loops may have physiological roles, the formation of stable, aberrant R-loops has been observed in disease, particularly neurological disorders and cancer. Despite the importance of these structures, methods to assess their distribution in the genome invariably rely on affinity purification, which requires large amounts of input material, is plagued by high level of noise, and is poorly suited to capture dynamic and unstable R-loops. Here, we present a new method that leverages the affinity of RNase H for DNA:RNA hybrids to target micrococcal nuclease to genomic sites that contain R-loops, which are subsequently cleaved, released, and sequenced. Our R-loop mapping method, MapR, is as specific as existing techniques, less prone to recover non-specific repetitive sequences, and more sensitive, allowing for genome-wide coverage with low input material and read numbers, in a fraction of the time.

## Main

R-loops are three-stranded nucleic acid structures that contain a DNA:RNA hybrid and a displaced single strand of DNA^1^. R-loops are dynamic structures whose levels are tightly controlled across the genome^2^^-^^4^. Alterations in nuclear R-loop levels are associated with disruption of transcription, DNA repair, and other key genomic processes ^5^^-^^9^. Identification of changes in R-loop abundance and distribution in different cell types could inform us on mechanisms that lead to cell type-specific pathology^10^^-^^15^. However, efforts to study the regulatory functions of R-loops have been hindered because of the sub-optimal methods used to enrich for and recover these chromatin structures. Therefore, there is a critical need to develop new methods that will allow for enhanced and systematic discovery of R-loops.

Currently, two distinct strategies are used to map the distribution of R-loops. The predominant strategy relies on the immunoprecipitation of chromatin containing R-loops using a monoclonal antibody, S9.6, thought to be specific for DNA:RNA hybrids^16^. DNA:RNA immunoprecipitation (DRIP) and all its variants ^17^^-^^20^ (bis-DRIP, S1-DRIP and RDIP), were foundational to the study of genome-wide R-loop localization, but share similar disadvantages: 1) they prepare chromatin for immunoprecipitation using harsh physical and biochemical treatments (high temperatures, strong detergents, sonication and/or prolonged enzymatic digestion of chromatin) in the absence of fixation, which might disrupt less stable R-loops before they can be detected, and 2) they rely on the S9.6 antibody whose strict specificity for DNA:RNA hybrids remains a subject of debate (e.g. it might also bind dsRNA^21^). The second, more recent, strategy to map R-loops takes advantage of the natural affinity of RNase H for DNA:RNA hybrids. RNase H is an enzyme that degrades the RNA strand of DNA:RNA heteroduplexes. Two published methods, DNA:RNA *in vitro* enrichment (DRIVE)^18^ and R-loop chromatin immunoprecipitation (R-ChIP)^22^ target R-loops by using a catalytic deficient version of RNase H (RHΔ) that retains its affinity for DNA:RNA hybrids but does not cleave the RNA strand. In both cases the DNA:RNA hybrids bound by RHΔ are enriched by affinity purification. In DRIVE, RHΔ is fused to the maltose-binding protein (MBP), incubated with sheared chromatin *in vitro* and bound R-loops recovered by affinity purification on amylose resin^18^. In R-ChIP, V5-RHΔ is expressed *in vivo*, and the R-loops recovered by immunoprecipitation using the V5 affinity tag^22^. Although both DRIVE and R-ChIP take advantage of the exquisite specificity of RNase H for targeting DNA:RNA species (as opposed to the more questionable specificity of the S9.6 antibody^21^) they still suffer from the limitations typical of affinity purifications, including high backgrounds, requirement for large amounts of starting material, and time consuming protocols.

To overcome these limitations, we have developed a new R-loop mapping strategy, termed “MapR”. MapR combines the specificity of RNase H for DNA:RNA hybrids with the sensitivity, speed, and convenience of the CUT&RUN approach^23,24^, whereby targeted genomic regions are released from the nucleus by micrococcal nuclease (MNase) and sequenced directly, without the need for affinity purification. Specifically, in MapR cells are immobilized (Fig. 1a, step 1) and permeabilized and a fusion protein comprising a catalytically inactive RNase H and MNase (RHΔ-MNase) is allowed to diffuse into the nuclei in absence of calcium ions, thus keeping the MNase enzyme inactive (Fig. 1a, step 2). After equilibration (Fig. 1a, step 3, top), calcium is added, and the nuclei incubated for 30 minutes at 0°C before stopping the reaction with EGTA. This results in the release of chromatin fragments targeted by RHΔ and therefore containing R-loops in their native state (Fig. 1a, step 4). As a control for MapR, we perform the same experiment using MNase lacking the RHΔ moiety (Fig. 1 step 3, bottom). Finally, released chromatin fragments are purified and sequenced (Fig. 1a, step 5).

**Figure 1:**
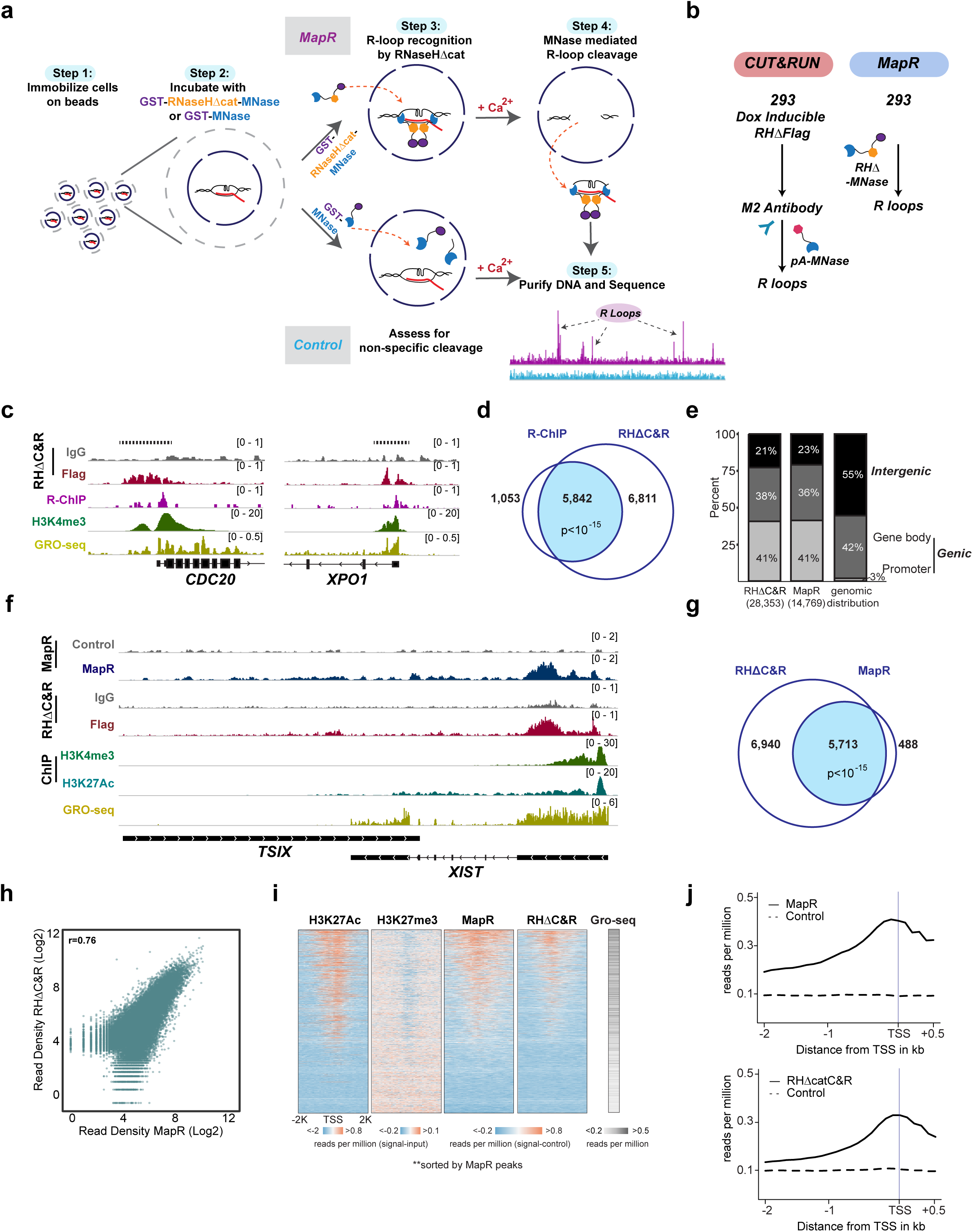
MapR, a native antibody-independent R-loop detection strategy. (a) R-loop recognition and recovery by MapR. Step 1: Cells are immobilized on concanavalin A beads and permeabilized. Step 2: Equimolar amounts of a catalytic deficient mutant of RNase H fused to micrococcal nuclease (GST-RHΔ-MNase) or GST-MNase is added to immobilized cells. Step 3: The RHΔ module recognizes and binds R-loops on chromatin. Step 4: Controlled activation of the MNase moiety by addition of calcium results in cleavage of DNA fragments in proximity to R-loops. Step 5: Released R-loops diffuse out of the cell; the DNA is recovered and sequenced. (b) Schematic of RHΔC&R using Flag M2 antibody (left) and MapR using GST-RHΔ-MNase (right) in HEK293. (c) Enriched regions identified by RHΔC&R and R-ChIP in HEK293. GRO-seq and H3K4me3 tracks indicate active gene transcription. The scale for the *y* axis is reads per million reads mapped (RPM). (d) Venn Diagram of gene level overlap between RHΔC&R and R-ChIP. Total number of unique genes with an R-loop at the promoter region (−2kb/+2kb from the TSS) and their overlap are shown. p<10^-15^, hypergeometric distribution. (e) Peak distribution of MapR and RHdeltaC&R showing percent of peaks mapping to promoter regions (−2kb/+2kb from the TSS), gene bodies (entirety of gene including introns, excluding promoter region), or intergenic regions. Total peak numbers are shown in parentheses. Background genomic distribution is shown for comparison. (f) MapR and RHΔC&R signals at the *XIST* and *TSIX* genes. GST-MNase and IgG controls are shown for MapR and RHΔC&R respectively. H3K4me3^36^ and H3K27Ac^31^ ChIP-seq, and GRO-seq^22^ tracks are shown as proxy for transcriptional activity. (g) Venn Diagram of gene-level overlap between RHΔC&R and MapR. Total number of unique genes with an R-loop at the promoter region (−2kb/+2kb from the TSS) and their overlap are shown. p<10^-15^, hypergeometric distribution. (h) Correlation scatterplot showing read densities for the union of peaks from MapR and RHΔC&R (Log2 scale). r= 0.76, Spearman correlation coefficient. (i) Heatmaps of H3K27Ac, H3K27me3, MapR and RHΔC&R signal intensity across all TSS sorted by MapR signal. GRO-seq signals were summed and collapsed into a box per gene. (j) Metagene plots of MapR (top) and RHΔC&R (bottom) signals at all TSSs.

As a first step toward developing MapR, we sought to determine whether a conventional, antibody mediated CUT&RUN approach in a cell line expressing tagged RHΔ could release R-loop-containing fragments. For this, we expressed a FLAG-tagged version of RHΔ containing a nuclear localization signal (Supplementary Fig. 1a) in HEK293 cells and performed a standard CUT&RUN assay using an anti-FLAG antibody to reveal the chromatin distribution of RHΔ, and therefore R-loops. (Fig. 1b, left). FLAG CUT&RUN for transgenic RHΔ (RHΔC&R) identified 28,353 peaks compared to an IgG control. These presumptive R-loops mapped to 12,653 genes, of which 5,842 overlapped with R-loop containing genes as identified by immunoprecipitation of RHΔ from crosslinked chromatin in the R-ChIP approach (Fig. 1c, 1d). This overlap is highly significant (p<10^-15^, hypergeometric distribution) indicating that CUT&RUN correctly recovers a large portion of previously identified R-loops. The majority of nuclear R-loops are known to occur co-transcriptionally ^18,25^. In agreement with this and consistent with R-ChIP, the majority (79%) of peaks identified by RHΔC&R occurred in genic regions, with 41% localized at promoters and 38% within the gene body (Fig. 1e). We conclude that R-loops can be targeted *in vivo* by RHΔ and their distribution can be revealed using a CUT&RUN approach.

Although the above strategy successfully retrieved native R-loops without affinity purification steps, it still required genetic manipulation of the cells to express a FLAG-tagged version of RHΔ. This presents obvious limitations when studying R-loops in cells that are difficult to transfect or sub-clone, such as, patient-derived primary cells that do not divide *in vitro.* To overcome these limitations, we reasoned that the FLAG antibody step could be bypassed by fusing RHΔ directly to MNase and providing this recombinant protein exogenously after cell immobilization and permeabilization. Toward this end, we expressed and purified GST-RHΔ-MNase (henceforth RHΔ-MNase) from *E.coli* (Supplementary Fig. 2a). As a control, we used GST-MNase in our experiments to assess for non-specific cleavage by MNase across the genome (Fig. 1a, Supplementary Fig. 2a). To ascertain that the presence of the RHΔ moiety did not affect the enzymatic activity of MNase, we digested chromatin with equimolar amounts of MNase and RHΔ-MNase. We found that the two fusion proteins had comparable enzymatic activity, since they produced similar patterns of nucleosomal ladders after 10 and 30 minutes (Supplementary Fig. 2b).

Next, we immobilized and permeabilized HEK293, and incubated them with either MNase (control) or RHΔ-MNase (MapR) (Fig. 1a). We activated the MNase moiety in both recombinant proteins by addition of calcium at the same time and for the same duration. As in CUT&RUN, we constructed libraries from cleaved DNA fragments that diffused out of the nucleus and sequenced them. Genome-wide profiles obtained by MapR (i.e. exogenous RHΔ-MNase fusion protein) were very similar to those obtained by expressing RHΔ *in vivo* and performing a conventional FLAG CUT&RUN (Fig. 1f), whereas no discernible signal was obtained with MNase alone. MapR enriched regions were predominantly genic (77%) with 41% pf peaks mapping to promoters and 36% within the gene body, consistent with the idea that this technology effectively identifies R-loops *in vivo* (Fig. 1e).

Co-transcriptional R-loops are known to occur at the 5’ end of active genes immediately downstream of the promoter and, to a lesser extent, at the 3’ end of active genes ^5,26,27^. As an example, we inspected the *XIST* long non-coding RNA (lncRNA) gene. HEK293 are female cells and therefore one of the two X chromosomes is subject to X chromosome inactivation, a process that is dependent on expression of the XIST lncRNA ^28,29^. Both MapR and RHΔC&R signals are clearly higher than the respective controls at the 5’ end of the *XIST* gene (Fig. 1f). *XIST* also contains an antisense gene, *TSIX* ^30^, that is expressed only in early development and is silent in HEK293. In contrast to the *XIST* locus, the *TSIX* gene showed no detectable signal from either MapR or RHΔC&R (Fig. 1f).

MapR using RHΔ-MNase identified 14,769 peaks compared to an MNase-only control (Fig. 1e). These peaks mapped to the promoters of 6,201 genes, of which 5,713 overlapped significantly (p<10^-15^) with promoter R-loop-containing genes as identified by RHΔC&R (Fig. 1g). Despite the ~7,000 genes where R-loop peaks were called by the peak-calling algorithm only in RHΔC&R, read densities from MapR and RHΔC&R over all the peaks were highly correlated (Fig. 1h, Spearman r=0.76), demonstrating that the two approaches detected broadly comparable genomic regions as being occupied by R-loops. We analyzed the strength of MapR and RHΔC&R signals (Fig. 1i) at all transcription start sites (TSS) and found that enriched regions from both datasets tracked closely with actively transcribed genes, as determined by GRO-seq, by the presence of the activating chromatin mark, histone H3 lysine 27 acetylation (H3K27ac)^31^, and by the corresponding depletion of the repressive chromatin mark H3K27 trimethylation (H3K27me3)^32^. No distinguishable MapR or RHΔC&R signal was observed at the TSS of inactive genes (Fig. 1i). Consistent with a predominant localization of R-loops near and upstream of TSSs, metagene analyses for both MapR and RHΔC&R revealed an accumulation of signal starting 2 kb upstream and peaking at the TSS (Fig. 1j). Thus, we conclude that genomic regions enriched by our MapR approach are specifically found at active genes and are broadly consistent with previously reported profiles for R-loops^22^. Importantly, these analyses show that MapR, a technique that bypasses the need for transgenic cells, identifies the same genomic regions as FLAG CUT&RUN performed on RHΔ-expressing cells.

Having demonstrated that regions identified by MapR have genomic features consistent with R-loops (i.e. they localized to the 5’ end of active genes), we next wished to determine if they also displayed known biochemical properties of R-loops. Bona fide R-loops are defined by the presence of a DNA:RNA heteroduplex, whose recognition by RNase H is the foundation for MapR. We reasoned that pre-treating immobilized and permeabilized cells with an enzymatically active RNase H would result in degradation of the RNA strand, restoration of double-stranded DNA, and loss of MapR signal (Fig. 2a). On the other hand, if our RHΔ-MNase fusion protein bound non-specifically to chromatin regions devoid of R-loops, these interactions should not be affected by pre-incubation with active RNase H. Indeed, MapR signal at the 5’ end of the *RWDD1* and *ANP32E* genes was considerably reduced by pre-treatment with active RNase H (Fig. 2b), an observation that held true throughout the genome (Fig. 2c, 2g), demonstrating that most if not all peaks detected by MapR contained DNA:RNA heteroduplexes. A similar RNase H-dependent reduction in signal intensities was observed in MapR experiments performed in U87T cell lines (Supplementary Fig. 3a and b).

**Figure 2:**
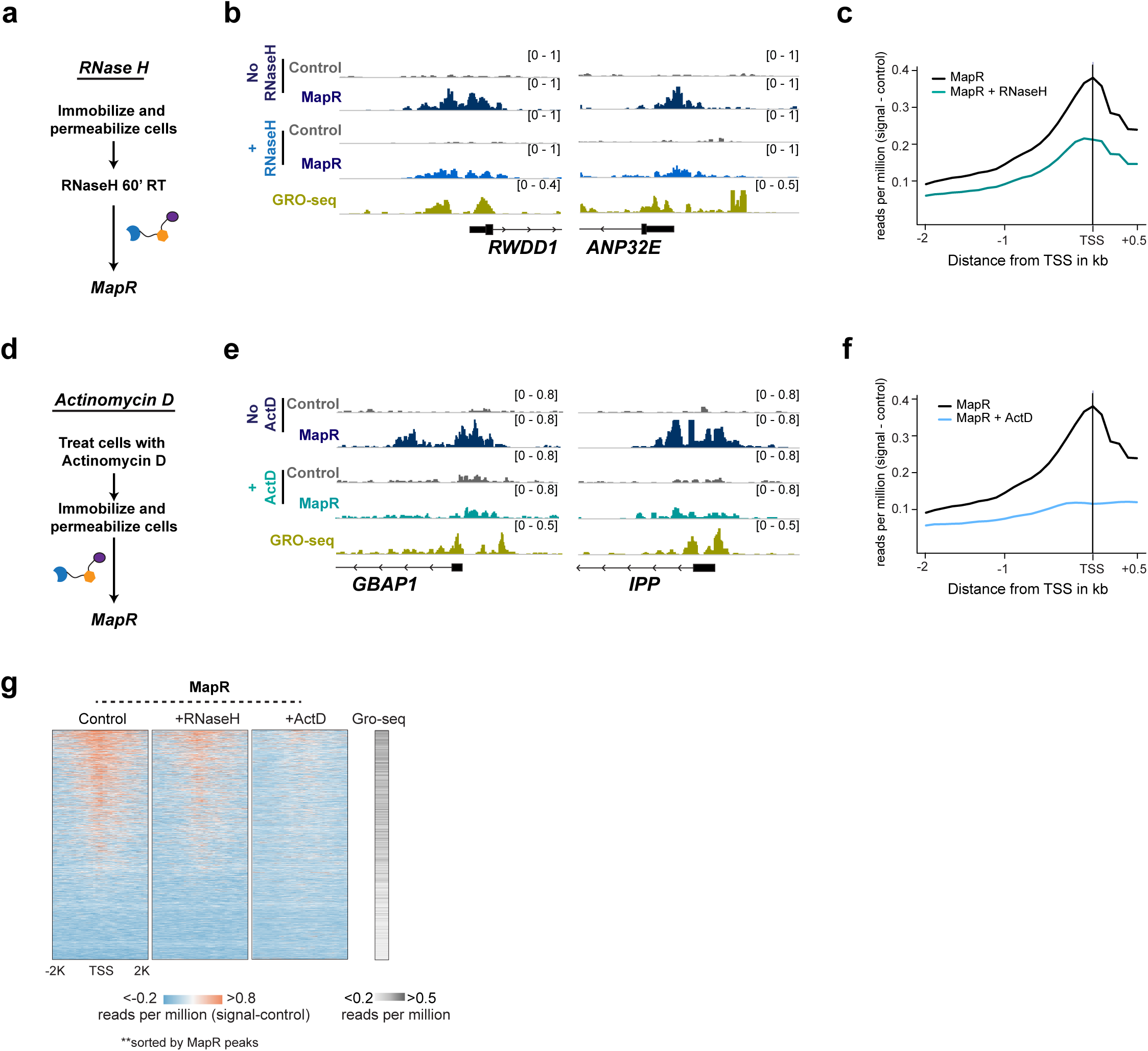
Characterization of R-loops obtained by MapR. (a) Schematic of RNase H treatment followed by MapR identification of R-loops. (b) Genome browser views *RWDD1* and *ANP32E* genes showing MapR signals with and without RNase H treatment. GRO-seq tracks show transcription at these genes. The scale for the *y* axis is in reads per million mapped (RPM). (c) Metagene plots of MapR signals at TSS of all genes with and without RNase H treatment. (d) Schematic of actinomycin D (ActD) treatment followed by MapR identification of R-loops. (e) Genome browser views of *GBAP1* and *IPP* genes showing MapR signals with and without ActD treatment. GRO-seq tracks show active transcription. (f) Metagene plots of MapR signals at TSS of all genes with and without ActD treatment. (g) Heatmaps of MapR signals across all TSS in control, RNase H and ActD treated HEK293 cells, sorted by MapR signal. GRO-seq signals from untreated HEK293 were summed and collapsed into a box per gene.

Since the majority of cellular R-loops are a consequence of active transcription, we reasoned that a general transcription inhibitor should cause decreased MapR signal (Fig. 2d). Consistent with this, treating cells with actinomycin D, an inhibitor of transcription elongation, caused a decrease in MapR signal at specific genes (*GBAP1* and *IPP*, Fig. 2e) and genome-wide (Fig. 2f and g; Supplementary Figs. 3c and d). These results show that the genomic regions recovered by MapR contain DNA:RNA hybrids that are degraded by RNaseH and whose formation is prevented by transcription inhibition. Therefore, MapR detects genomic features with the biochemical properties of R-loops *in vivo*.

Next, we asked how MapR compared to existing R-loop detection strategies. We selected for comparison the two methods representing the two strategies outlined above: RDIP for methods that use the S9.6 antibody to purify DNA:RNA hybrids and R-ChIP for methods that employ RNase H. Importantly, datasets obtained with these techniques in the same cell type (HEK293) were publicly available ^19,22^. Visual inspection of the genome browser revealed that the MapR signal broadly resembled that of RDIP and R-ChIP (Fig. 3a). Promoters that contained an R-loop according to MapR overlapped significantly (p<10^-15^) with genes identified by R-ChIP or RDIP (Fig. 3b); however, MapR detected thousands of additional genes as compared to both previous technologies (Fig. 3b), raising the question of whether these newly detected genes contained bona fide R-loops and were previously missed.

**Figure 3:**
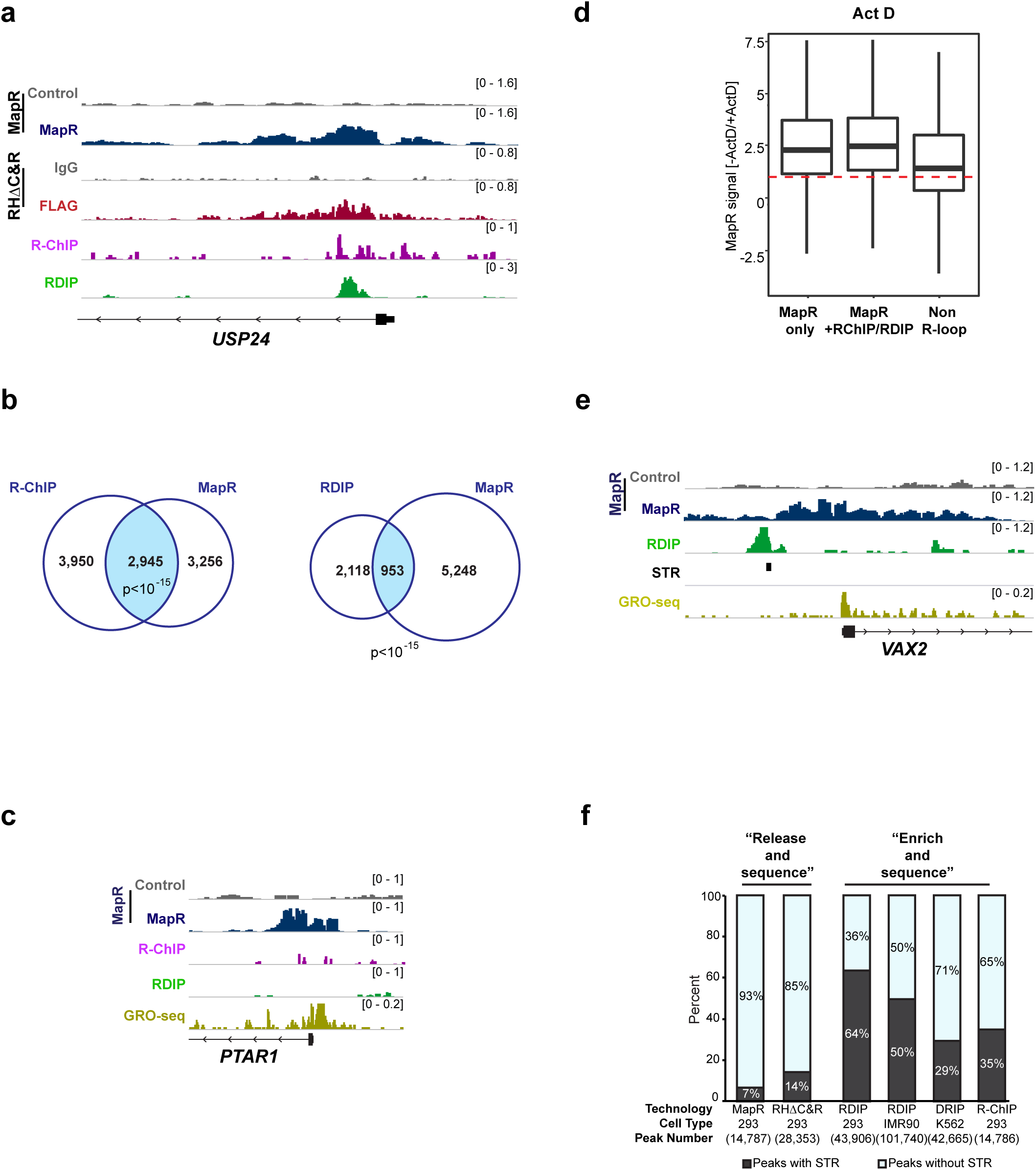
Similarities and differences between MapR and other R-loop detection methods. (a) Genome browser view of the *USP24* gene showing MapR, RHΔC&R, RDIP and R-ChIP signals. The scale for the *y* axis is in reads per million mapped (RPM). (b) Gene-level overlap between MapR and R-ChIP datasets (left) and MapR and RDIP datasets (right). Total number of unique genes with an R-loop at the promoter region (−2kb/+2kb from the TSS) and their overlap are shown. p<10^-15^, hypergeometric distribution (c) Genome browser view of *PTAR1* that shows MapR but no R-ChIP or RDIP signals. GRO-seq tracks indicate active transcriptional status. (d) Ratio in the −2kb/+2 kb window around the TSS in MapR signal in untreated cells and cells treated with Actinomycin D to inhibit transcription. Genes were divided in three classes: genes containing an R-loop only according to MapR (left), genes containing an R-loop according to MapR as well as R-ChIP or RDIP (middle), and genes that did not contain R-loops. (e) Genome browser view of *VAX2*, that shows MapR signals that do not overlap with RDIP peaks in close proximity. Simple tandem repeat (STR) and GRO-seq tracks are shown. The scale for the *y* axis is in reads per million mapped (RPM). (f) Percent of peaks that contain STRs in MapR, RHΔC&R, RDIP and R-ChIP experiments in HEK293. Results from published RDIP (IMR90)^19^ and DRIP (K562)^25^ datasets are also shown.

To evaluate whether the signals obtained exclusively from MapR experiments correspond to genuine R-loops, we first visually inspected these regions to determine whether they correspond to regions of active transcription, as ascertained by the presence of a GRO-Seq signal (Fig. 3c; Supplementary Figs. 4a and 4b). Next, we compared the effect of actinomycin D treatment (see Figs. 2d–g) on presumptive R-loops detected exclusively by MapR with those in common between the techniques. Actinomycin D treatment resulted in a similar or increased reduction of MapR signal in MapR-only genes compared to genes in common between MapR and R-ChIP/RDIP (Fig. 3d and Supplementary Fig. 4c). The MapR signal over a control gene set that according to all techniques did not contain R-loops did not show any appreciable change upon actinomycin D, confirming treatment specificity. Finally, we analyzed the distribution of sequences predicted to give rise to G-quadruplex structures, which is a common feature of the displaced DNA strand in R-loops^18,22,33-35^. The frequency of G-quadruplexes in the promoter regions of genes with promoter R-loops by MapR were comparable as those measured in genes called using our RHΔC&R as well as the other two R-loop detection strategies, while non R-loop genes have a lower frequency of G-quadruplexes in their promoters (Supplementary Fig. 4d). Thus, MapR identifies bona fide native R-loops.

We further assessed the sequencing depth required in our experiments compared to other R-loop detection methods (R-ChIP^22^ and RDIP^19,22^). We analyzed all datasets from 293 cells using decreasing amount of reads to ask if some techniques required fewer reads to achieve sensitive R-loop detection. As compared to R-ChIP and RDIP, MapR and RHΔC&R showed clearly enriched regions with lower read numbers (Supplementary Figs. 5a and 5b), which is consistent with the advantage that a CUT&RUN approach confers over conventional ChIP or other affinity enrichment methods^23,24^.

We further analyzed the published data and observed that R-ChIP peaks had a genomic distribution similar to MapR, with a majority of peaks (79%) mapping to genes and a small number (21%) to intergenic sites. In comparison, 49% of RDIP peaks occurred at intergenic sites and only 51% within genes (Supplementary Fig. 6a). Of the genic peaks only 8% mapped to promoter regions in the RDIP dataset. While investigating this discrepancy, we noticed that RDIP peaks were frequently proximal to but not overlapping with MapR peaks and that these sites of exclusive RDIP enrichment often overlapped with simple tandem repeats (STRs) (Fig. 3e and supplementary Fig. 6b). In HEK293 cells, we found that 64% of RDIP peaks contained STRs, whereas only 7% of MapR peaks contained STRs (Fig. 3f). To determine whether this STR enrichment was observed in other technologies, we analyzed RHΔC&R and R-ChIP from HEK293. We also analyzed published RDIP and DRIP datasets from IMR90 and K562 cells respectively to exclude experimental- and cell type-specific bias^19,25^. We found that MapR and RHΔC&R, which rely on cleavage and release of nucleic acid followed by direct sequencing (as opposed to the enrichment strategies used in RDIP, DRIP, and R-ChIP) showed lower overlap with STRs (Fig. 3f). Interestingly, peaks called by RDIP showed a frequency of STR overlap that correlated with the strength of peak enrichment (Supplementary Fig. 6c). Such a correlation was absent in the MapR, RHΔC&R, R-ChIP, and DRIP datasets.

Finally, we wished to probe the detection limits of our system. Our MapR experiments above were performed with five million cells, which is within the range used in most ChIP and DRIP experiments; however, a main advantage of CUT&RUN over immunoprecipitation methods to map chromatin marks (ChIP) is that the lack of an affinity purification step largely decreases the amount of input material required^23,24^. Thus, we tested whether MapR could identify R-loops starting from 50-fold fewer cells. The MapR profiles obtained from 10^5^ cells closely resembled those obtained with 5 million cells (supplementary Fig. 6d), with a similar genome-wide enrichment at and upstream of active TSSs (Supplementary Figs. 6e and 6f). Thus, we conclude that MapR offers the ability to discover R-loops with high sensitivity and is robust even when cell numbers are limiting. Notably, these improvements on sensitivity and specificity are accompanied by a greatly streamlined experimental protocol that can be completed in 1 day, which is ~4X less than the fastest alternative.

In summary, MapR is an efficient, convenient, and fast method to generate genome-wide maps of R-loops. MapR employs an antibody independent strategy that can be used in any cell type without the need to generate stable transgenic lines. Importantly, MapR can identify R-loops in small cell numbers which can facilitate its future application to study aberrant R-loops formed in diseases using patient-derived material.

## Supplementary Figure legends

**Supplementary Figure 1:**
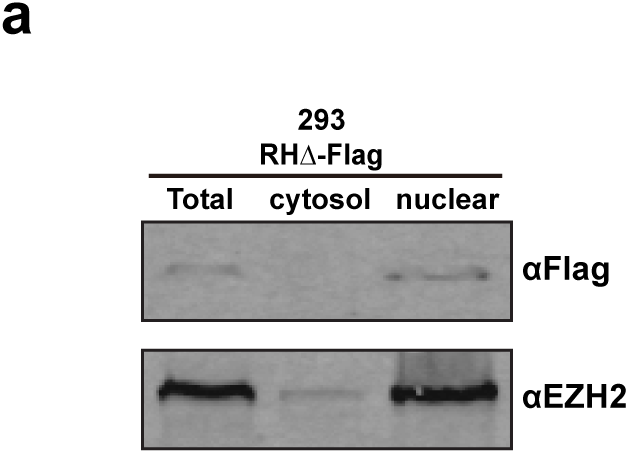
Expression of RHΔ protein in HEK293. Western blot for RHΔ-Flag protein in nuclear and cytosolic fractions using anti FLAG antibodies. EZH2, a nuclear protein, was used as a control for fractionation.

**Supplementary Figure 2:**
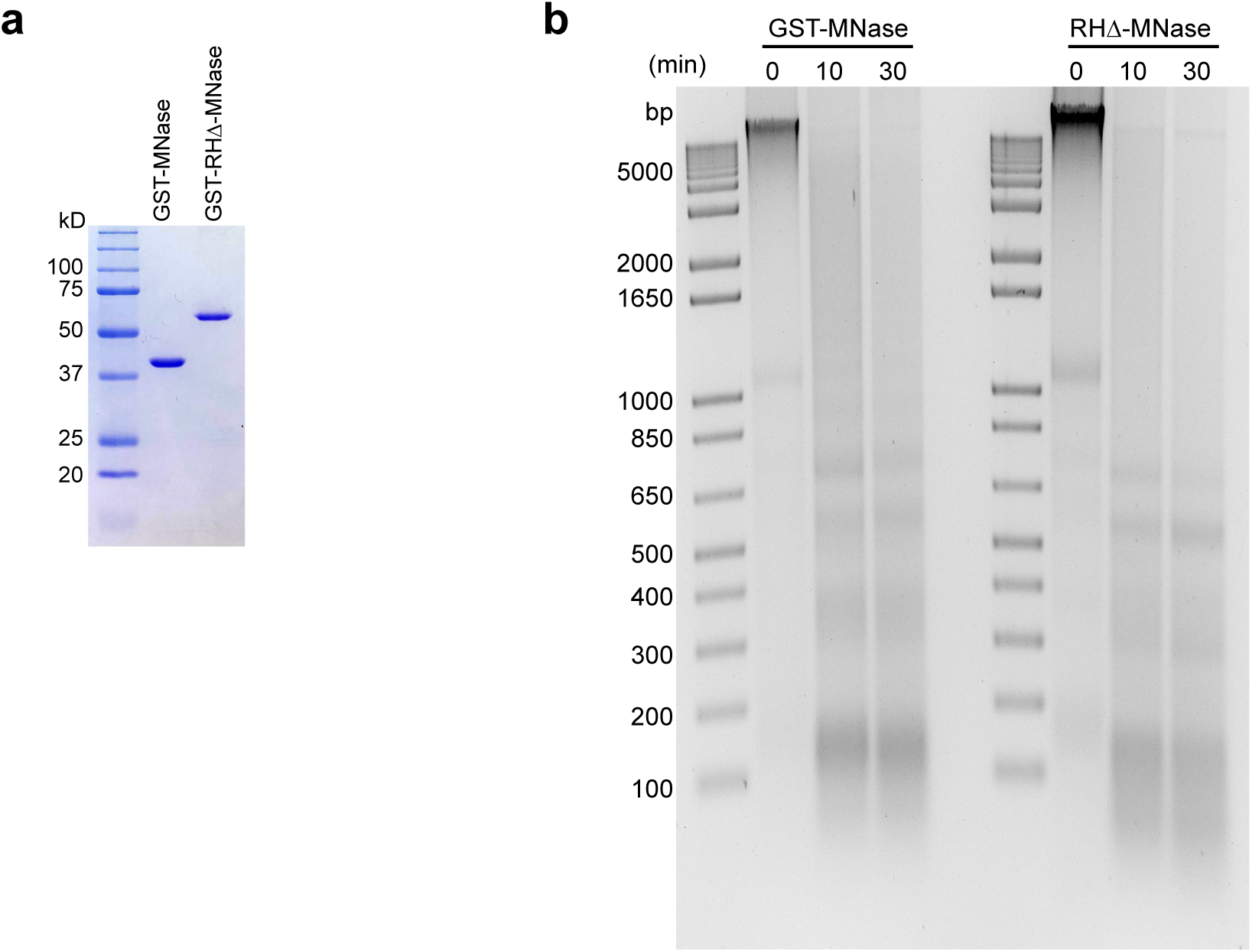
Purification and characterization of GST-MNase and GST-RHΔ-MNase proteins. (a) Coomassie blue stain of purified GST-MNase and GST-RHΔ-MNase proteins. (b) Chromatin digestion assay using equimolar amounts of GST-MNase and GST-RHΔ-MNase proteins at different time-points as indicated above the gel. Purified DNA from digestion reactions was resolved on a 2% agarose gel and stained with ethidium bromide.

**Supplementary Figure 3:**
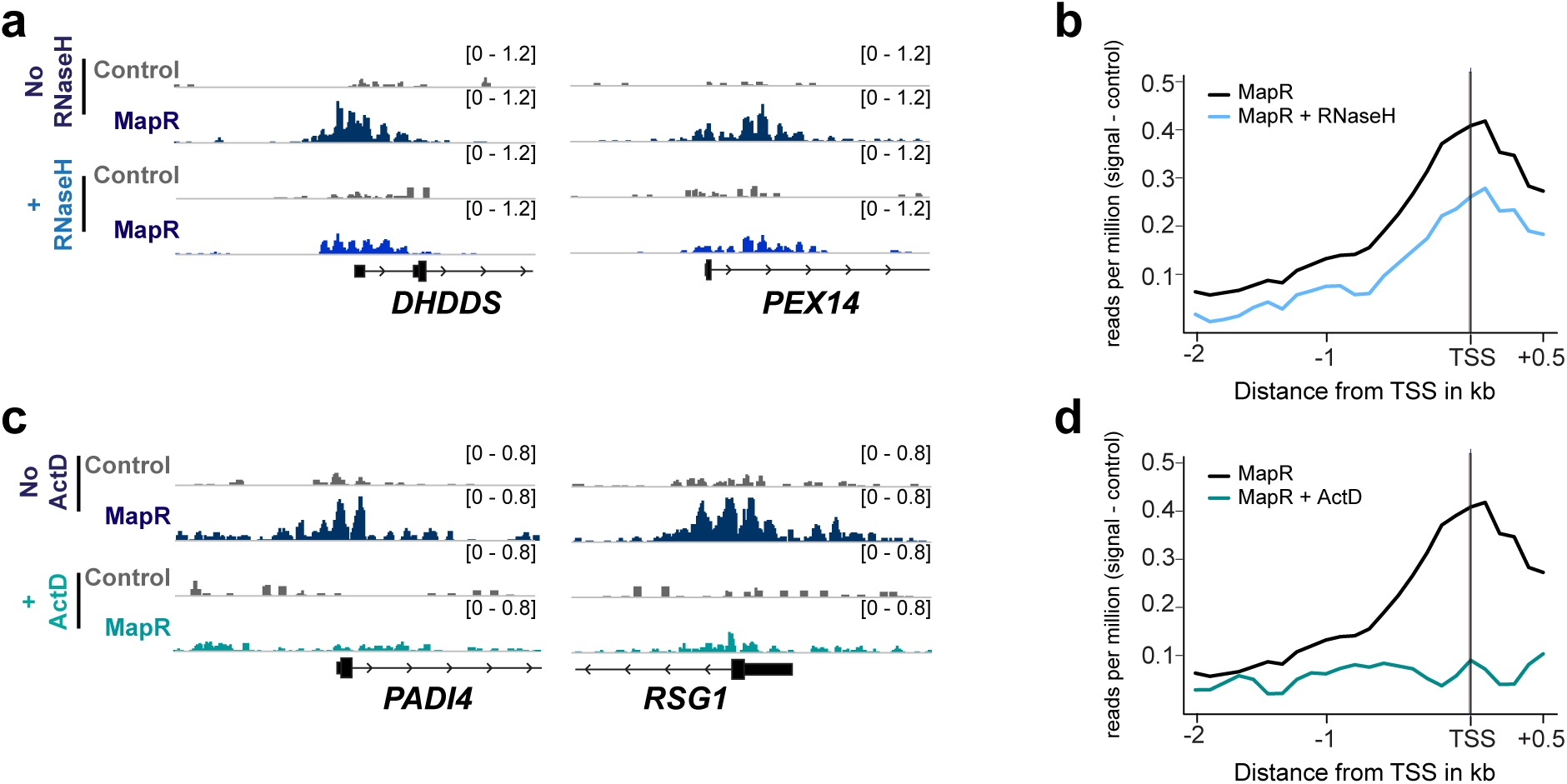
MapR enriched regions in U87T cells are bona fide R-loops. (a) Genome browser views of *DHDDS* and *PEX14 genes* showing MapR signals with and without RNase H treatment. The scale for the *y* axis is in reads per million mapped (RPM). (b) Metagene plots of MapR signals in U87T cells at TSS of all genes with and without RNase H treatment. (c) Genome browser views of *PADI4* and *RSG1* genes showing MapR signals in U87T cells with and without ActD treatment. The scale for the *y* axis is in reads per million mapped (RPM). (d) Metagene plots of MapR signals in U87T cells at TSS of all genes with and without ActD treatment. (e) Heatmaps of MapR signals across all TSS in control, RNase H and ActD treated U87T cells, sorted by MapR signals. GRO-seq signals were summed and collapsed into a box per gene.

**Supplementary Figure 4:**
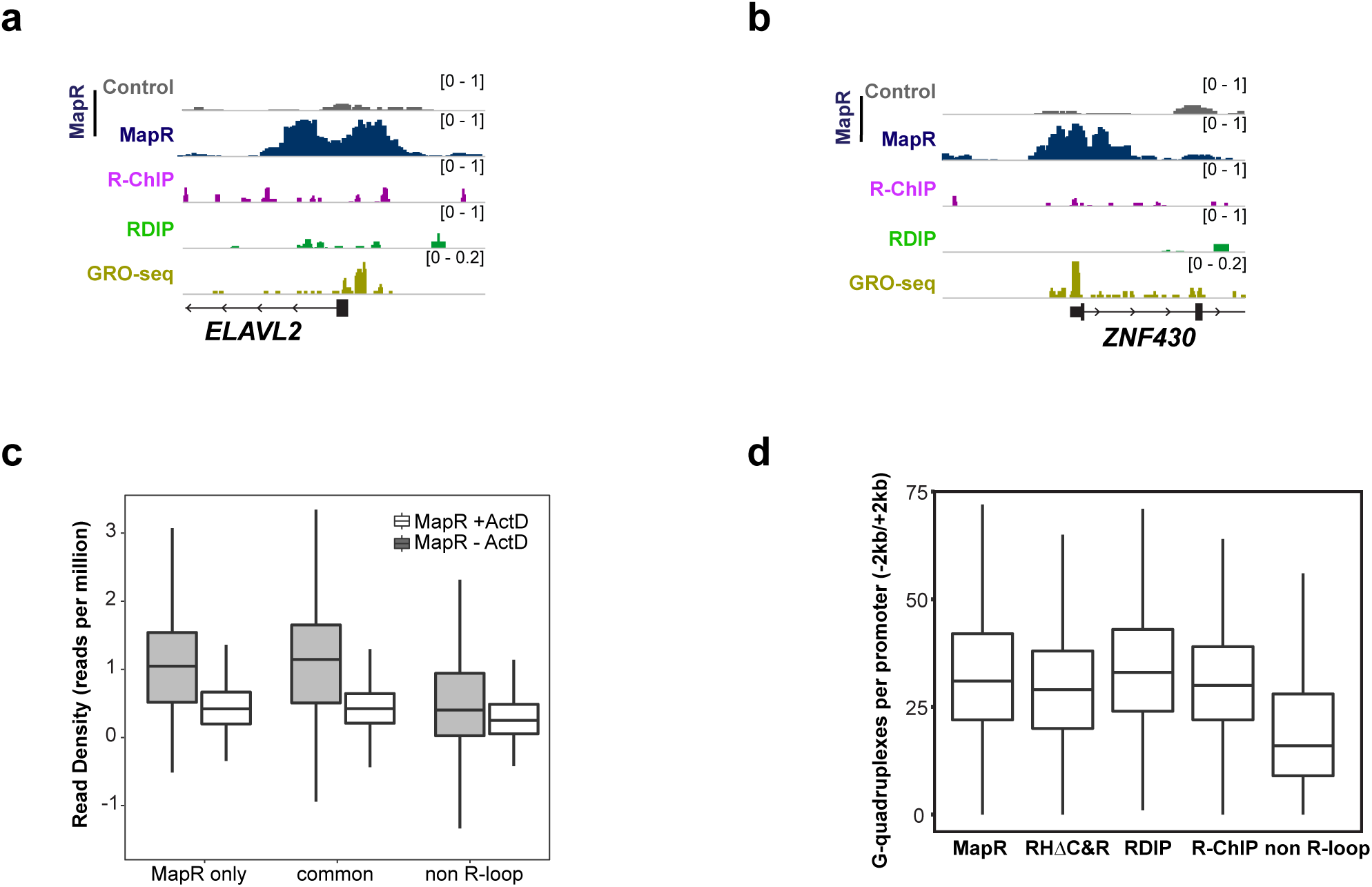
R-loops exclusively identified by MapR show reduced MapR signals upon treatment with actinomycin D. (a) Genome browser view of *ELAVL2*, that shows MapR but no R-ChIP or RDIP signals. GRO-seq tracks indicate active transcriptional status. (b) Genome browser view of *ZNF430*, that shows MapR but no R-ChIP or RDIP signals. GRO-seq tracks indicate active transcriptional status. (c) MapR signal upon Actinomycin D treatment across genes identified by MapR only, by MapR and R-ChIP or RDIP and genes that did not contain any R-loops. Read densities are sum of read per million (RPM) in the promoter region (−2kb/+2kb). (d) Genes called from MapR, RHΔC&R, R-ChIP and RDIP in 293 cells show similar frequency of predicted G quadruplex structures at their promoter, indicative of R-loop presence, while non R-loop genes have a lower frequency of G quadruplexes.

**Supplementary Figure 5:**
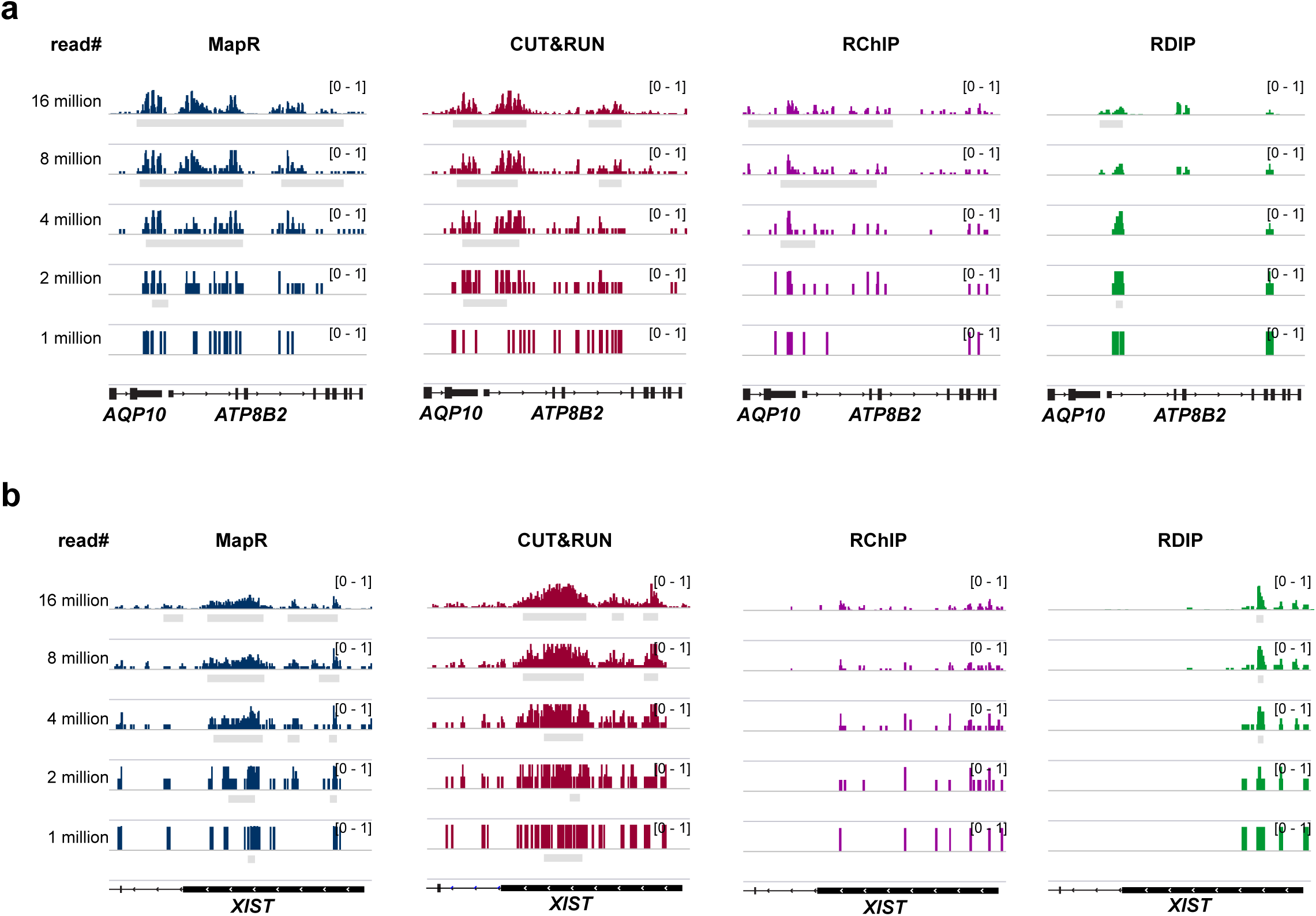
Prediction of G quadruplex structures in different R-loop detection strategies. (a,b) Genome browser views of *ATP8B2 and XIST*. Replicates and their controls from each technology were downsampled to the indicated number of reads. Peaks were called using the same parameters as for full samples. The scale for the *y* axis is in reads per million mapped (RPM).

**Supplementary Figure 6:**
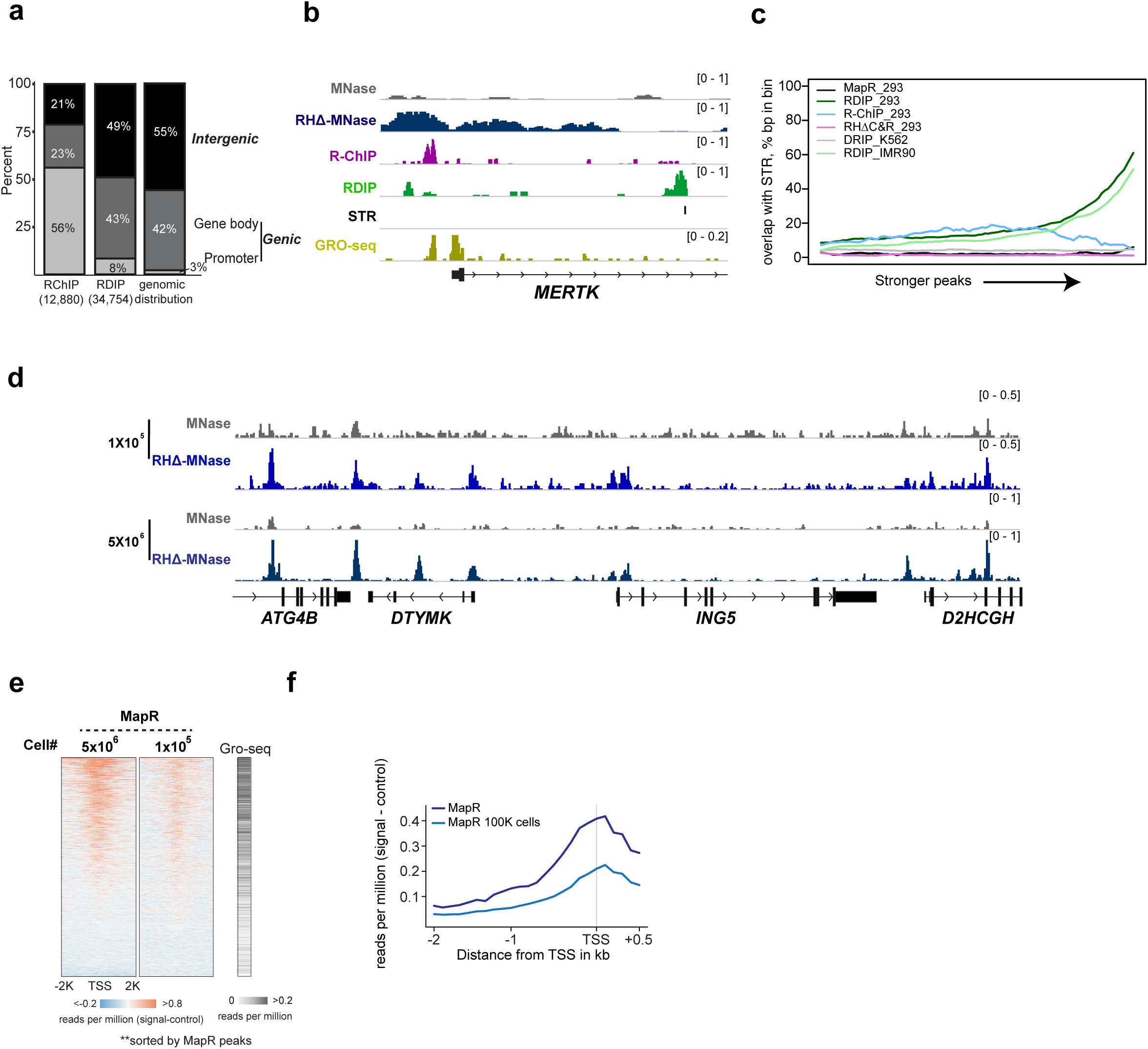
Analysis of simple tandem repeat enrichment across R-loop detection methods. (a) Peak distribution of R-ChIP and RDIP showing percent of peaks mapping to promoter regions (−2kb/+2kb from the TSS), gene bodies (entirety of gene including introns, excluding promoter region), or intergenic regions. Total peak numbers are shown in parentheses. Background genomic distribution is shown for comparison. (b) Genome browser view of *MERTK*, that shows overlapping MapR and R-ChIP signals that do not overlap with RDIP peaks in close proximity. Simple tandem repeat (STR) and GRO-seq tracks are shown. (c) Line plot showing % of peaks (by bp), binned by q-value, that overlap STRs. (d) Genome browser view showing MapR signals obtained from 1X10^5^ and 5X10^6^ cells. (e) Heatmaps of MapR signals at TSS of all genes in 1X10^5^ and 5X10^6^ cells. (f) Metagene plot of MapR signals at TSS of all genes in 1X10^5^ and 5X10^6^ cells.

**Supplementary Table 1.**
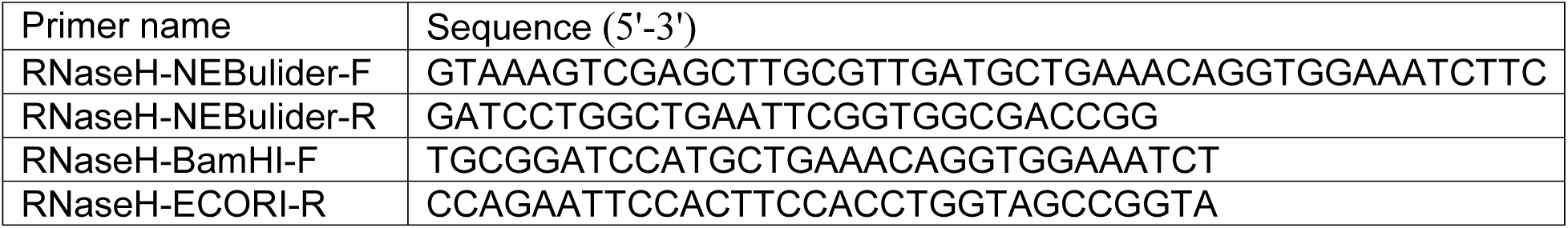

## Materials and Methods

### Plasmid construction

RNaseHdcat was amplified from pICE-RNaseHI-D10R-E48R-NLS-mCherry (Addgene plasmid: 60367) and sub-cloned into pGEX-6p-1-MNase and pLT3GEPIR^37^. Primer sequences can be found in supplementary table 1.

### Cell culture

HEK293 and U87T cell lines were grown in DMEM supplemented with 10% serum. Stable cell lines were generated by transfection with Lipofectamine 2000 (Invitrogen) and selection with puromycin (1µg/ml). Protein expression was induced by addition of doxycycline (1µg/ml final concentration) and analyzed by western blot with antibodies as indicated.

### Protein expression and purification

GST-MNase and GST-RHΔMNase were expressed in BL21(DE3) (ThermoFisher) using standard expression conditions and purified using GST-agarose beads (Affymetrix) as per manufacturer’s instructions. Purified proteins were stored in BC100 buffer (25 mM Tris-HCl pH7.6, 0.2 mM EDTA, 100mM KCl, 1 mM β-mercaptoethanol) containing 25% glycerol.

### MNase activity assay

3 × 10^6^ HEK293 cells were resuspended in Buffer A (10 mM MES pH 6.5, 0.25M Sucrose, 60 mM KCl,15 mM NaCl, 5 mM MgCl_2_, 1 mM CaCl_2_, 0.5% Triton X-100, 0.5 mM PMSF) and incubated on ice for 20 min. Cells were centrifuged at 1000g for 10 min and resuspended in 160 µl Buffer B (10 mM PIPES pH 6.8, 50 mM NaCl, 5 mM MgCl_2_, 1 mM CaCl_2_, 0.1 mM PMSF) and divided into two tubes. 1.5µM of GST-MNase and GST-RHΔ-MNase proteins was added and chromatin digestion performed at 37°C . 25 µl of the digestion reaction was transferred at different time points (0, 10, 30 min) to a tube containing 1µl 0.5 M EDTA, 15 µl 10% SDS, 10 µl 5 M NaCl and 40 µl H_2_O. DNA was extracted using phenol-chloroform and resolved on 2% agarose gels.

### MapR and CUT&RUN

CUT&RUN was performed exactly as described in (REF) using 5µg of FLAG M2 antibody or mouse IgG. MapR buffer volumes and incubation times follow the standard CUT&RUN protocol unless otherwise specified. 5×10^6^ cells were washed with twice with 1.5 ml room temperature wash buffer (20 mM HEPES pH 7.5, 0.15 M NaCl, 0.5 mM Spermidine, 1 mM protease inhibitors) and immobilized on Cocanavalin A-coated beads. Immobilized cells were divided equally into two tubes and resuspended in 50 µl wash buffer (20 mM HEPES pH 7.5, 0.15 M NaCl, 0.5 mM Spermidine, 1 mM protease inhibitor) containing 0.02% Digitonin. GST-MNase and GST-RHΔ-MNase proteins were added to a final concentration of 1µM and incubated overnight at 4 °C with rotation. The beads were washed three times, resuspended in 100 µl Dig-wash buffer and place on ice. 2 µl 0.1 M CaCl2 added to activate MNase and digestion was carried out for 30 minutes. Reaction was stopped by adding equal volume of 2x STOP buffer (340 mM NaCl, 20 mM EDTA, 4 mM EGTA, 0.02% Digitonin, 5µg RNaseA, 5 µg linear acrylamide and 2 pg/ml heterologous spike-in DNA). The samples were incubated at 37 °C for 10 minutes to release the protein-DNA fragments and spun down at 16000g for 5 minutes at 4 °C. Supernatants were transferred to fresh tubes, 2 µl 10% SDS and 5 µg proteinase K was added and reactions were incubated at 70 °C for 10 minutes. DNA was extracted using phenol-chloroform.

For RNase H treatment, after immobilization of cells to beads, 150 U RNase H in 50 µl Dig-wash buffer was added and incubated at room temperature for 1 hr before proceeding with MapR. HEK293 and U87T cells were treated with Actinomycin D (5µg/ml) for 8 hrs and processed for MapR.

### Library preparation and sequencing

DNA was end-repaired using End-It Repair Kit, tailed with an A using Klenow exo minus, and ligated to custom adapters with T4 DNA ligase. Fragments > 150 bp were size-selected with SPRI and subjected to ligation-mediated PCR amplification (LM-PCR) with custom barcoded adapters for Illumina sequencing using Q5 DNA polymerase. All enzymes except Q5 (NEB) were from Enzymatics (a Qiagen company). Sequencing was performed on a NextSeq 500 (Illumina).

### Sequencing analysis

Raw reads were mapped to the human genome (hg19) with Bowtie2^38^ with default parameters. Normalized genome-wide read densities were computed using deeptools ^39^. Peaks were called for each sample (with associated control as background, if possible) using MACS 2.1.1 ^40^ using the parameters: --broad --broad-cutoff 0.1.

Peak locations were computed by identifying R-loops in promoter regions, (−2kb/+2kb of the TSS), gene bodies (the entirety of the gene including introns, but excluding the promoter region), and intergenic regions. Gene level overlaps were calculated by identifying genes in the hg19 NCBI RefSeq gene set with an R-loop at the promoter for each technology and reporting common genes. GRO-seq raw data was downloaded from GSE97072 (293 cells) and GSE92375 (U87 cells). Peaks were called using the HOMER tool findPeaks^41^. Any gene with overlapping Gro-seq peak(s) was considered active, while genes without GRO-seq peaks were considered inactive. Heatmaps were created using pheatmap. Reduction of signal with ActD treatment was calculated using the total occupancy across the window from −2kb to +2 kb of each unique TSS. Read densities were computed (bedtools coverage) over the merged peak co-ordinates from MapR and RHΔC&R and normalized to total mapped reads for each dataset.

### STRs

Short tandem repeats in hg19 were downloaded from UCSC. For percent of peaks overlapping with STRs, a peak was considered to contain an STR if it had at least a 1 bp overlap with an annotated STR. For the peak strength analysis, peaks were binned by q-value into 100 bins of even size. For each bin, the % of bp of all peaks in the bin that overlapped with an STR was reported.

### G-quadruplexes

G-quadruplexes were detected in promoters (−2kb/+2kb of TSS) of genes with an R-loop in the promoter region, as well as genes with no promoter R-loop, for all technologies using pqsfinder ^42^ and the number of G-quadruplexes per promoter was reported.

### Published data

RDIP data (293, K562, IMR90) were downloaded from GEO: GSE68948. R-ChIP and Gro-seq data were downloaded from GEO: GSE97072. H3K27me3 ChIP-seq data were downloaded from GEO: GSM855015. H3K27ac ChIP-seq data were downloaded from ENCODE: ENCSR000FCH. U87 Gro-seq data were downloaded from GSE92375.

### Data availability

Sequencing data generated for this study have been deposited in the NCBI GEO as GSE120637. Data will remain private during peer review and released upon publication.

## Acknowledgments

We thank Guohong Li (IBP,Chinese Academy of Sciences) for pGEX-6p-1-MNase plasmid and Hongwu Zheng (Cornell) for U87T cells. This work was supported by the NIH New Innovator Award DP2-NS105576 to K.S. R.B. acknowledges financial support from the NIH (R01GM127408) and the Searle Scholars Program (15-SSP-102). E.S. acknowledges financial support from the NIH (T32HG000046).

